# Reinforcement Failing guides the discovery of emergent physical dynamics in adaptive tumor therapy

**DOI:** 10.1101/2025.04.08.647768

**Authors:** Serhii Aif, Maximillian Eiche, Nico Appold, Elias Fischer, Timon Citak, Jona Kayser

## Abstract

Artificial intelligence is revolutionizing scientific discovery in medicine, with reinforcement learning (RL) emerging as a promising tool for optimizing therapeutic strategies. Yet applying RL to complex scenarios such as therapy dynamics in solid tumors is constrained by the challenge of constructing training environments that are both computationally efficient and mechanistically interpretable. Here we introduce Reinforcement Failing, an AI-guided, human-in-the-loop discovery framework that shifts the focus from agent policy optimization to the refinement of the training environment itself. By combining multi-fidelity RL with group-relative performance evaluation across agent cohorts, Reinforcement Failing systematically reveals emergent mechanisms that first-principles models overlook. We apply this framework to adaptive therapy in solid tumors, which seeks to delay resistance-mediated treatment failure. In this setting, Reinforcement Failing uncovered a coupling between the mechanically driven collective motion of cells and spatially-heterogeneous proliferation that strongly influences therapy outcomes. Incorporating these emergent physical mechanisms into an augmented training environment improved cross-environment therapeutic performance and exposed potential pitfalls in translation. More broadly, these findings position Reinforcement Failing as a powerful artificial scientific discovery framework, capable of deciphering high-complexity processes at the interface of physics, machine learning, and medicine.

## Main

Despite substantial advances in modern cancer treatments — including targeted therapies and immunotherapies — the emergence of therapy resistance remains one of the most urgent challenges in modern oncology, calling for innovative approaches [1]. While many patients initially respond to antitumor therapy, long-term tumor control often remains elusive due to the presence of resistant subpopulations. Building on mathematical models of tumor dynamics, preclinical and clinical pilot studies of evolution-based adaptive therapies (AT) have recently demonstrated how intratumor competition between susceptible and resistant subpopulations may be exploited to extend the time to disease progression (Fig. 1 a) [2–11].

**Figure 1:**
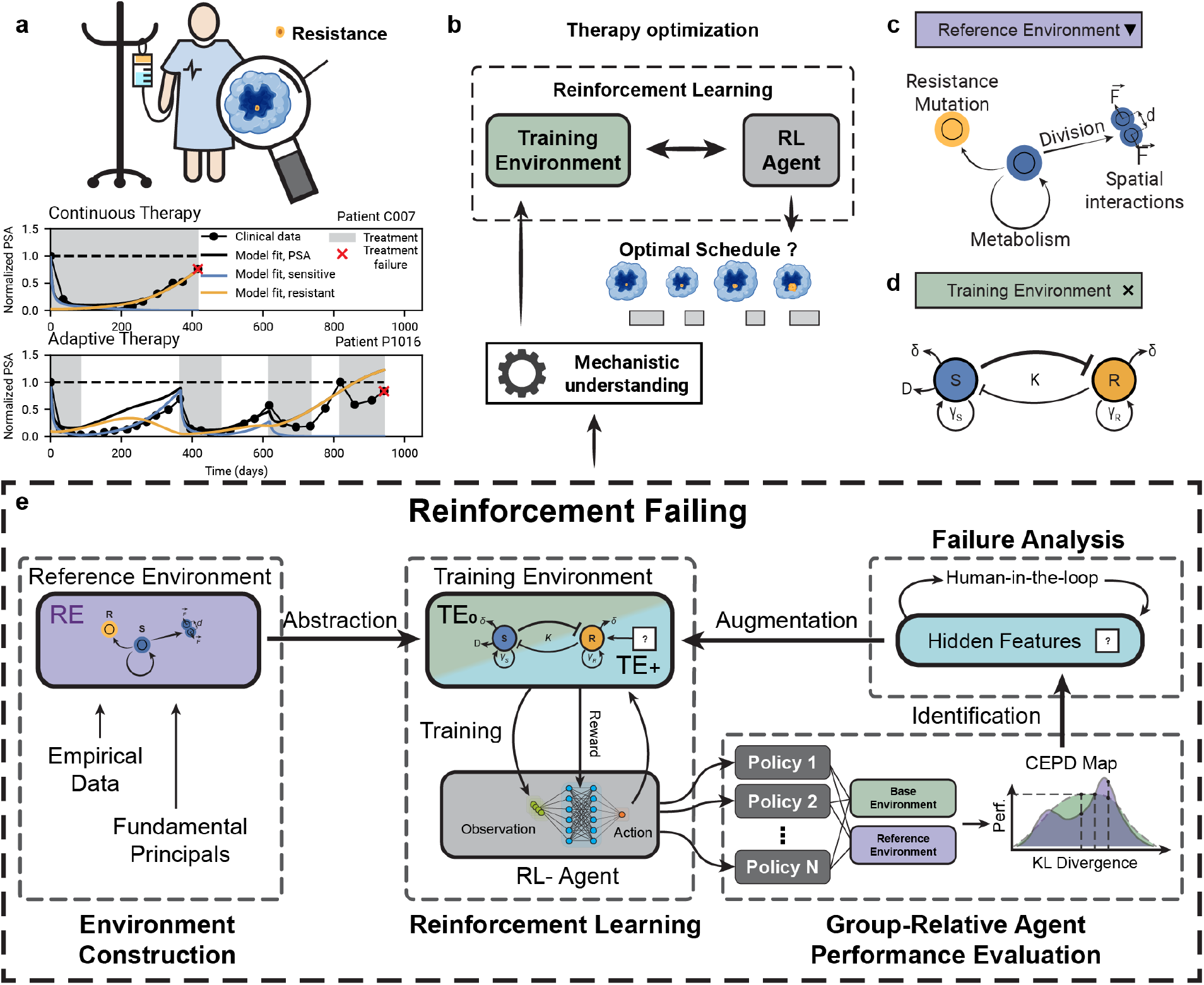
Overview of the Reinforcement Failing framework. **a** Illustration of a cancer patient with a tumor that has developed resistance. Prostate cancer patients data - normalized prostate-specific antigen (PSA) values for continuous and adaptive therapy AT_50_ (adapted from Ref. [3]). Solid lines represent a mathematical model fit to individual patient data as reported in that study. **b** Schematic of therapy optimization via Reinforcement Learning using a mechanistically interpretable training environment. **c** Schematic of the reference environment, explicitly simulating the growth, division, mutation, and force-induced mutations of thousands of cells. **d** Schematic of mathematical tumor growth model based on coupled ordinary differential equations, used as the training environment (see main text for details). **e** Overview of the Reinforcement Failing framework, consisting of four main blocks. Environment Construction: Based on empirical data and fundamental physical principles, a high-fidelity simulation is designed as reference environment (RE). The complex reference environment is then abstracted into a mathematical base training environment (TE_0_). Reinforcement Learning: An ensemble of *N* independent agents is trained in TE_0_. Group-Relative Agent Performance Evaluation (GRAPE): Evaluating all agents in both TE_0_ and RE yields a cross-environment performance difference (CEPD) map. Failure Analysis: CEPD-guided analysis of differentially performing agents by a human researcher enables the identification of performance-critical but previously hidden system features and their integration into a refined augmented training environment (TE_+_). The augmentation is followed by the training and evaluation of second-generation agents. Tumor schematic created in BioRender. Kayser, J. (2025) https://BioRender.com/3jkoj49.

The transition from fixed dosing regiments toward adaptive, evolution-informed strategies reflects a broader shift in medicine toward flexible, datadriven interventions — a shift increasingly enabled by advances in artificial intelligence [12]. Within ongoing efforts to optimize dosing in chemo- and radiotherapy [13–19], recent studies by Gallagher *et al*. [20] and Lu *et al*. [21] have applied reinforcement learning (RL) directly to adaptive therapy optimization via more nuanced dosing strategies (Fig. 1b).

Since training of RL agents is typically performed in simulations, a central challenge lies in identifying which system characteristics must be captured by these training environments to ensure transferability of the obtained policy to the original context. In packed cellular aggregates, including solid tumors, this “abstraction” is particularly challenging due to the intricate coupling of spatial population structure, resource gradients, and mechanical cell–cell interactions, all of which can drive collective phenomena that alter evolutionary trajectories and treatment responses [22–25]. Computational and experimental studies have shown that these mechanisms strongly influence resistance evolution and the potential benefits of adaptive dose scheduling [26–34].

Modeling such processes explicitly makes simulations both opaque for scientific interpretation and computationally prohibitive for RL. The nascent field of multi-fidelity RL aims to address these challenges by pairing low-fidelity training environments with subsequent evaluation and possibly transfer learning in high-fidelity simulations [35, 36]. However, without principled guidance on which emergent physical mechanisms shape therapeutic outcomes, this abstraction process becomes a bottleneck, potentially omitting key dynamics [37].

To overcome this critical hurdle, this work introduces a novel framework that integrates high-fidelity simulations of physical tumor growth dynamics with group RL to guide the systematic augmentation of the low-fidelity training environment (Fig. 1c-d). The core idea behind this approach is “Reinforcement Failing” (RF) (Fig. 1e), a novel concept that leverages Group-Relative Agent Performance Evaluation (GRAPE) to expose hidden system features that shape therapeutic success. Instead of discarding the policies of presumably “failing” RL agents, we show that these can serve as informative signal for human researches, guiding them toward hidden emergent processes that differentiate between therapy success and failure. Using this RL-guided human-in-the-loop approach, we uncover that position-dependent growth and mechanically-induced cell motion are key drivers of therapy outcomes. Augmenting the training environment accordingly yields superior performing agents while facilitating a detailed understanding of the underlying mechanisms and ensuing therapeutic pitfalls by human researchers. In the following, we first outline the general architecture of the framework before applying it to the specific scenario of adaptive tumor therapy.

## Results

### Reinforcement Failing: A framework for RL-guided augmentation of abstracted training environments

The Reinforcement Failing framework consists of four sequential main blocks: construction of the reference environment and its abstraction into a training environment, training of agent ensemble via reinforcement learning, cross-environment group-relative agent performance evaluation, and the human-in-the-loop augmentation of the training environment by previously hidden features (Fig. 1 e).

The initial step is the construction of the highfidelity reference environment (RE), designed to capture the intricacies of the complex real system, followed by its subsequent abstraction into a low-fidelity but high-efficiency base training environment (TE_0_). The base training environment is then used to independently train an ensemble of treatment-decision-making agents with a diverse range of policies via reinforcement learning.

A key innovation of the framework is the use of Group-Relative Agent Performance Evaluation (GRAPE) to uncover hidden but outcome-critical emergent features. Rather than aiming to find the single best policy, GRAPE moves beyond absolute performance metrics as the central goal but instead aims to systematically explore the decision space more broadly. To this end, each agent is evaluated in both the training and reference environments to generate a cross-environment performance difference (CEPD) map of the policy space (see Methods for details).

Using the Kullback–Leibler divergence between state–action probability distributions as a distance metric to hierarchically order policies in behavior space reveals the most informative agents and guides human researchers in identifying key behavioral patterns underlying performance mismatches. Understanding these failure modes provides the foundation for augmenting the training environment with the newly identified features. The augmented environment (TE_+_) is then used to train a cohort of second-generation agents, which are subsequently analyzed with GRAPE. If necessary, this cycle can be repeated for iterative model refinement until performance in the training and reference environments converges.

### Bridging tumor physics and adaptive therapy via Reinforcement Learning

To assess the role of emergent physics-driven dynamics in a therapy setting, we construct a high-fidelity reference environment (RE) based on the state-of-the-art PhysiCell cancer simulator [38–42]. In this model, individual cells are represented as mechanically interacting spheres that move freely in continuous space in response to external forces (Fig. 1c and Fig. 2). Cells proliferate by consuming diffusible resources and can be killed upon drug exposure. Any higher-order population phenomena, such as competition, is solely mediated via the abundance of local resources (Supplementary Movie 1).

**Figure 2:**
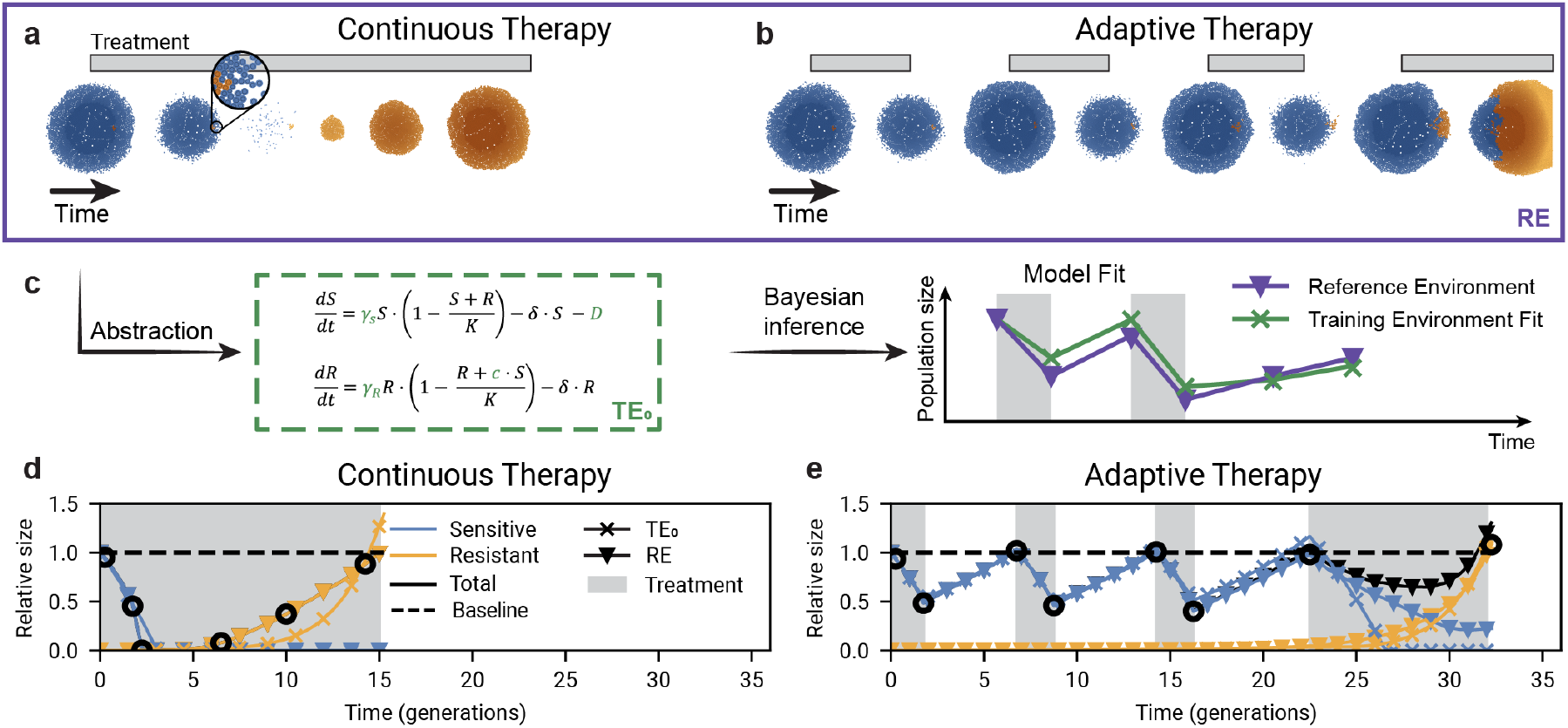
A cell-based tumor simulation as reference environment integrates emergent physical effects and therapy dynamics. **a, b** Snapshots of the RE simulation under Continuous Therapy (CT, Supplementary Movie 2, **a**) and Adaptive Therapy (AT50, Supplementary Movie 3, **b**). Sensitive cells are shown in blue, resistant cells in orange; brighter colors indicate higher proliferation rates. Gray bars indicate treatment application. **c** Schematic of the abstraction of reference environment dynamics into a fitted training environment. **d, e** Dynamics of sensitive (blue) and resistant (orange) population sizes, relative to the total initial population, are shown for RE (triangles) and the fitted TE_0_ (crosses), under CT (**d**) and AT50 (**e**). Grey regions denote treatment application, and black circles mark the time points shown in the snapshots.

The RE provides a detailed representation of emergent system characteristics, but its computational demands limit its suitability for training. Therefore, we first abstract the high-fidelity physics-based RE into a more tractable, low-fidelity mathematical training environment (Fig. 2c).

As initial base training environment (TE_0_), we use a system of coupled ordinary differential equations that describe the competition between drug-sensitive (*S*) and drug-resistant (*R*) populations (Fig. 1d), following previous work [20, 43]:

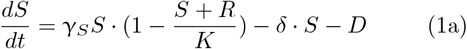

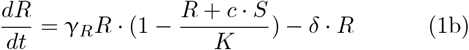

The environment models logistic growth of the two competing populations, with growth rates γ_*S*_ and γ_*R*_, limited by a shared carrying capacity *K* and an asymmetric competition coefficient *c*. Cell death is modeled through a stochastic rate δ, and the susceptible population additionally experiences a drug-induced death *D*, which is set to zero in the absence of treatment. In contrast to previous implementations, the drug effect is modeled independently of population size to account for the resource-limited periphery-to-core killing observed in the reference environment (see Methods for details).

We evaluated two treatment protocols in the RE: continuous therapy (CT) (Supplementary Movie 2) and adaptive therapy (AT_50_) (Supplementary Movie 3) (Fig. 2 d,e). The adaptive therapy protocol is analogous to the design of the clinical study by Zhang *et al*., pausing treatment when the tumor burden falls below 50% of its baseline value and resuming treatment when the burden returns to 100% [2, 3]. The dynamics observed in our *in silico* model qualitatively capture the trajectories reported in the clinical trial, including an extended control of tumor growth (compare Fig. 1 a). This is an inherent behavior of the system without any fitted parameters. Notably, the observed benefit of adaptive scheduling is not contingent on a resistance-associated fitness costs, corroborating previous reports on adaptive therapy in lattice-based spatial models [43, 44]. The parameters of the base training environment (TE_0_) are fitted to the dynamics under both strategies observed in RE via Bayesian inference (see Methods for details).

While it is clear that the simplified TE_0_ may not capture the whole spectrum of system dynamics, we want to test how well the complex behavior of the physical reference environment can be abstracted into *effective* parameters. For example, in the reference environment, many cells are confined to the resource-depleted tumor core, resulting in a lower effective proliferation rate γ_*S*_ than the actual division rate in the physical simulation. This abstraction process is analogous to the fitting of mathematical models to clinical patient data (Fig. 1a, b) [2, 3, 20, 30]. Note, these effective parameters inferred from this simplification should not be interpreted as reflecting the true biological rates in the system.

Despite its inherent limitations, the parameterized training environment qualitatively captures the trajectories of the reference environment (Fig. 2 a). To quantitatively compare strategy performance, we measure the time to therapy failure (TTF), defined as the time at which resistant cells reach the initial tumor size, rendering continued confinement elusive (see Methods).

We find that AT_50_ significantly outperforms CT, approximately doubling TTF in both the mathematical and the physical environments (CT: 15.3 *±* 0.2 generations in TE_0_ and 14.9 *±* 0.5 generations in RE; AT_50_: 33.8 ± 0.8 generations in TE_0_ and 30.2 ± 2.9 generations in RE; Extended Data Fig. 1). Based on these observations, we form the null-hypothesis that TE_0_ sufficiently captures the dynamics of the RE for reinforcement learning.

### Application of Reinforcement Failing to adaptive tumor therapy

To apply Reinforcement Failing in adaptive tumor therapy, we trained a cohort of *N* = 10 independent agents in the parameterized base training environment TE_0_ to explore the therapy decision space under the given model assumptions (Fig. 3a). Agents were trained via proximal policy optimization (PPO) and a customized reward function, using a sliding window of only the total normalized tumor burden as observation (see Methods for details). During training, all agents successfully converged to effective policies (Fig. 3 b). By learning how to use adaptive treatment application to optimally balance overall population growth and the risk of competitive release of resistant cells, trained agents achieved an average TTF of 149 ± 19 generations in the training environment (Fig. 3c). This represents an almost fivefold improvement over the fixed-threshold AT_50_ strategy (33.8 ± 0.8 generations).

**Figure 3:**
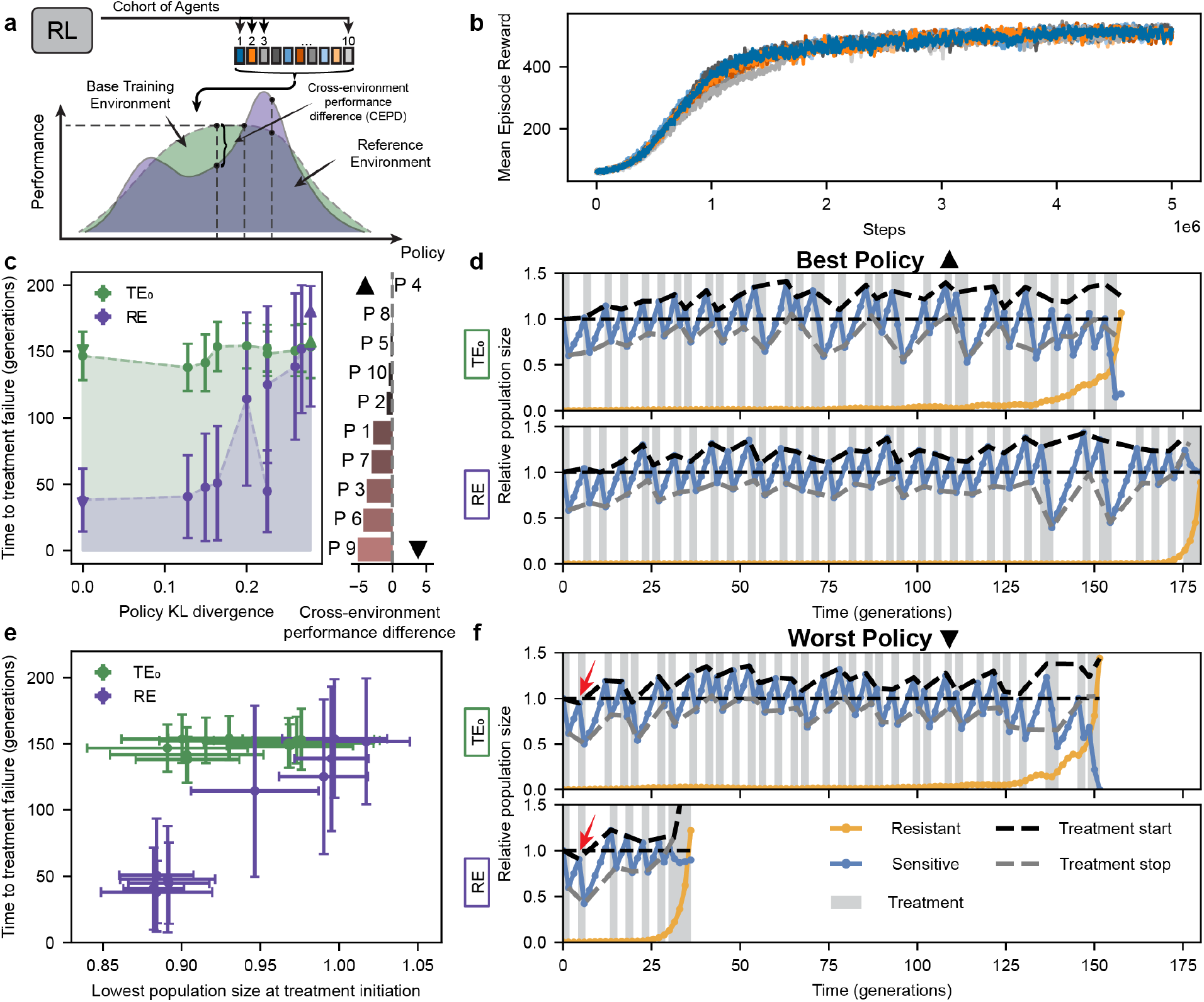
Reinforcement learning and GRAPE reveal critical agent behavior. **a** Schematic of agent policy space exploration and cross-environmental performance difference. **b** Mean episode reward over training steps of *N* = 10 trained agents in the TE_0_. **c** Left: Time to treatment failure (TTF) in TE_0_ (green) and RE (purple) for policies trained in TE_0_ (*N* = 10). Dots represent averages over *n* = 50 runs, error bars indicate the standard deviation. The Kullback-Leibler divergence of each policy from the worst-performing policy is used as behavior space coordinate. Right: Standardized cross-environment performance differences (Cohen’s d) are shown for all (*N* = 10) policies. Identified best (**▲**) and worst (**▼**) policies are indicated. **d, f** Examples of the dynamics of the sensitive (blue) and resistant (orange) relative population sizes for the best and worst performing policies in the reference environment, shown for both TE_0_ and RE (Supplementary Movies 4 and 5). Shown runs are indicated in **c** with **▲** (best) and **▼** (worst), colored according to the environment. Grey regions indicate treatment application. Dashed curves show the tumor burdens at treatment initiation (black) and treatment pause (gray). Arrows point to the application of treatment at low tumor burden by the agent using the worst policy (see main text). **e** Time to therapy failure (TTF) in TE_0_ (green) and RE (purple) for *N* = 10 policies trained in TE_0_ as a function of the lowest total population size at which treatment was initiated. Dots represent averages over *n* = 50 runs. Error bars indicate standard deviation.

However, applying the GRAPE evaluation framework reveals a substantial heterogeneity in agent performance when evaluated in the reference environment (Fig. 3c; Extended Data Fig. 2). This lack of transferability despite successful training rejects our null-hypothesis and instead suggests that the TE_0_ base training environment is insufficient.

To understand cause of the observed performance heterogeneity, we analyze the decision-making behavior and environment-state trajectories of poorly performing agents and compare them to those of agents that generalize well (Figs. 3d-f). We find that a key distinction lies in their treatment initiation patterns. High-performing agents initiate treatment at progressively higher tumor burdens, delaying aggressive therapy until tumor regrowth accelerates. In contrast, poorly performing agents have at least one early decision in which they apply treatment when tumor sizes are already reduced (see arrows in Fig. 3f). This episode is of little consequence in the training environment as long as it is followed by sufficient tumor regrowth (Fig. 3f, top). However, in the reference environment, once a sufficiently sharp reduction in tumor burden occurs, the system is unable to recover, resulting in uncontrolled growth of resistant cells and ultimately therapy failure (Fig. 3f, bottom).

To quantitatively test whether aggressive treatment at low tumor burdens is a predictor of agent failure, we measure the smallest tumor size at which each agent initiates treatment and analyze its correlation to the time of therapy failure (Fig. 3e). No significant correlation is observed in the training environment (Pearson’s correlation coefficient, *r* = 0.45). In contrast, we find a strong correlation (*r* = 0.98) in the reference environment, with well- and poorly-performing agents forming distinct clusters. Conversely, we find that a high average total burden the conventionally proposed driver of resistance confinement - strongly correlated with long TTF in the training environment (*r* = 0.91) but was a much weaker predictor in the reference environment (*r* = 0.77) (Extended Data Fig. 3)[44, 45]. These observations suggest additional mechanisms as determinants of therapy failure dynamics (see Supplementary Information and Extended Data Fig. 4 for a detailed analysis).

### Growth-driven cell motion accelerates resistance escape

Having pinpointed the agent behaviors responsible for divergent performance, we can now investigate the underlying mechanisms by analyzing the full spatiotemporal dynamics of relevant subtrajectories in a human-in-the-loop manner (Fig. 4a; Supplementary Movie 6).

**Figure 4:**
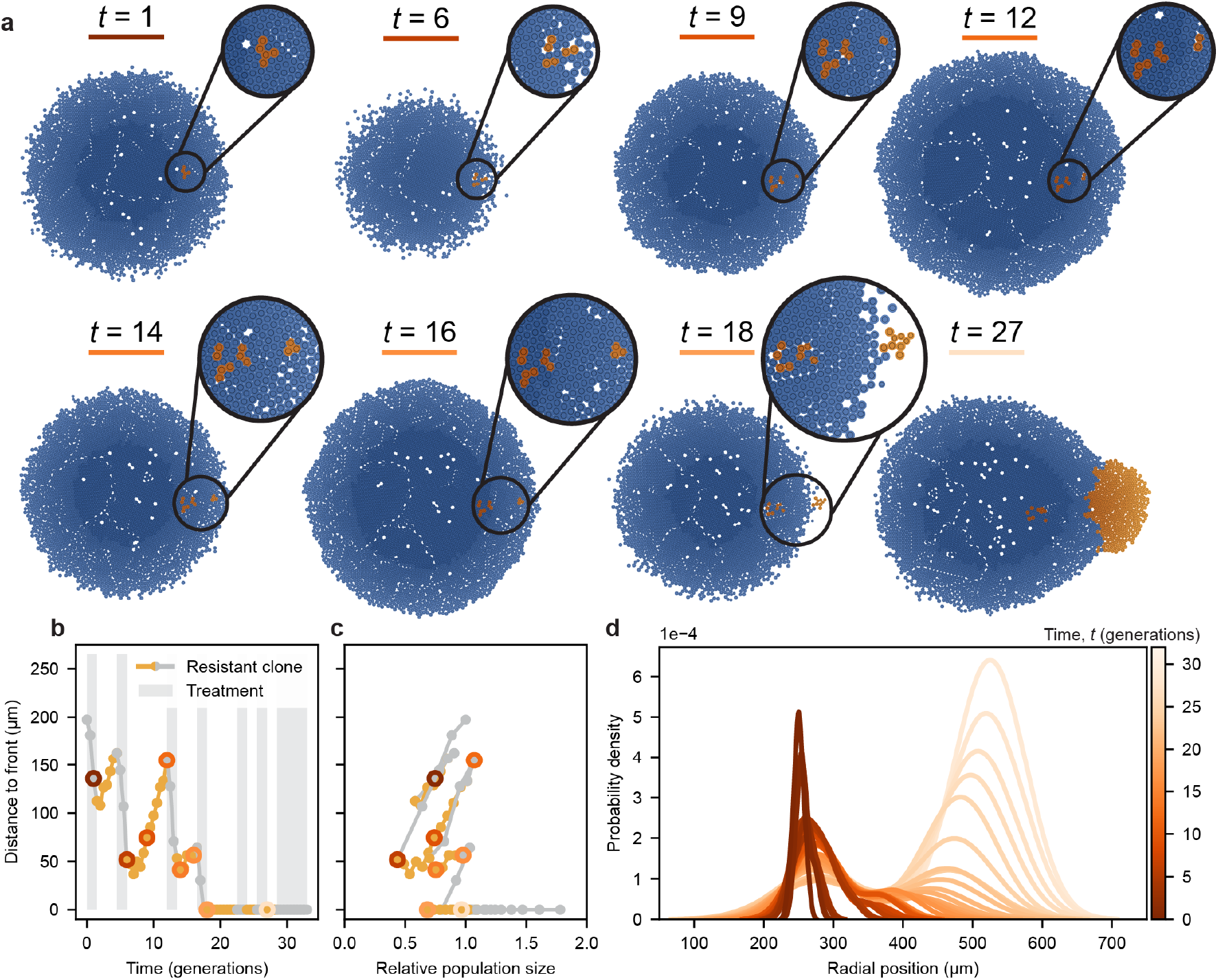
A mechanical ratchet effect drives the competitive release of resistant cells. **a** Selected simulation snapshots of the RE under the worst-performing policy. Sensitive cells in blue; resistant cells in orange. Brighter colors indicate larger growth rates (see Supplementary Movie 6). **b** Closest distance from the outer-most resistant cell to the nearest point on the tumor front over time for the simulation run in **a**. Orange lines indicate treatment holidays, gray indicates that treatment was applied. **c** Relationship between the distance to the front and the total population size, relative to the initial tumor burden. Colored circles around markers indicate the time points corresponding to the snapshots in **a. d** Radial distribution of resistant cells in time for the same simulation run. Brighter shades represent later time points.

We find that during the post-treatment expansion phase, resistant cells within the bulk can be physically displaced by the collective growth dynamics following a therapy cycle, especially if preceded by a substantial reduction of the susceptible population

(Fig. 4a). Importantly, once the distance of a resistant cell to the front is sufficiently decreased, even less intensive subsequent therapy cycles continue this process, gradually shifting the cell closer to the tumor front (Fig. 4b, c). This ratchet-like motion progresses in a stepwise but irreversible fashion until resistant cells reach the edge. As their distance to the tumor front decreases, resistant cells experience more favorable growth conditions and begin to expand more rapidly (Fig. 4d).

This interplay between position-dependent growth and mechanically induced motion of resistant cells is not captured by the base training environment. A refinement of the training environment is necessary to enable RL agents to account for these effects.

### Spatial augmentation of the training environment allows agents to avoid hidden therapy pitfalls

To gradually augment the training environment with the most essential spatial dynamics first, we focus on updating the representation of the resistant population while keeping the growth equation for susceptible cells unchanged (Fig. 5a). Two primary candidates for environment augmentation, suggested by the observed ratchet-like motion of resistant cells, are spatially heterogeneous proliferation and growth-induced pushing of cells.

**Figure 5:**
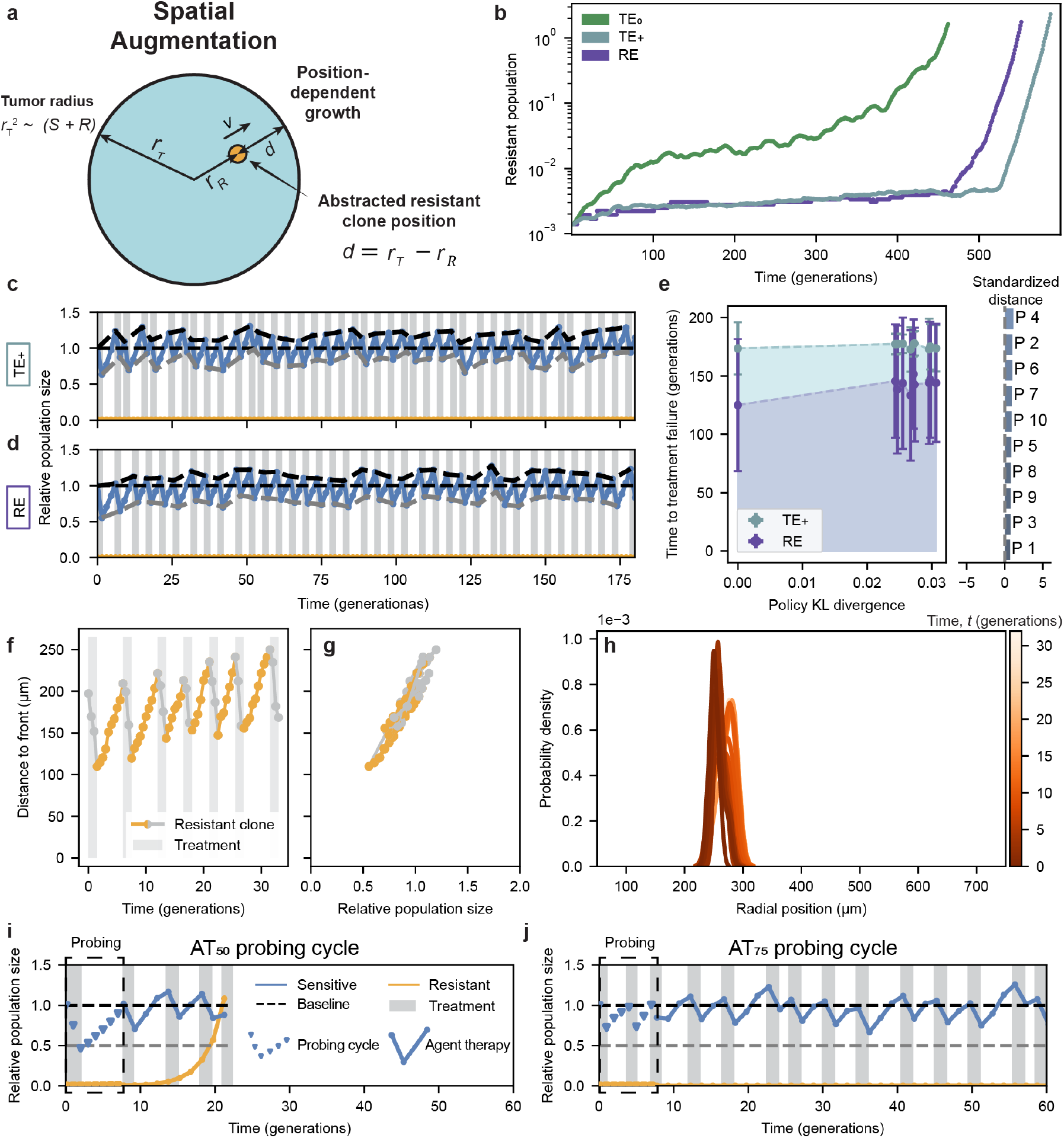
Augmenting the training environment to reflect spatial dynamics yields superior agent performance. **a** Schematic of the augmented environment (TE_+_). **b** Resistant population dynamics over time in the three environments, TE_0_ (green), TE_+_, (gray-blue), and RE (purple). **c, d** Relative population size dynamics of sensitive (blue) and resistant (orange) cells for a median-performing agent trained in the augmented environment evaluated in both TE_+_ and RE, respectively (Supplementary Movie 7). Gray bars indicate treatment application. Dashed curves show the tumor burdens at treatment initiation (black) and treatment pause (gray). **e** Time to treatment failure (TTF) in TE_+_ (gray-blue) and RE (purple) for policies trained in TE_+_ (*N* = 10). Dots represent averages over *n* = 50 runs, error bars indicate the standard deviation. Policies are spaced on the horizontal axis based on their Kullback-Leibler divergence, using the poorest-performing policy as reference. **f** Closest distance from outer-most resistant cell to the nearest point on the tumor front over time for the simulation run shown in **d**. gray indicates treatment application, orange indicates treatment holidays. **g** Distance of resistant clone to the front as a function of total population size. **h** Radial distribution of resistant cells in time for the same simulation run. Brighter shades represent later time points. **i, j** Dynamics of sensitive and resistant populations under the probing cycles for 8 generations of AT_50_ and AT_75_ therapies, respectively, followed by treatment decisions of the best performing agent in TE_+_.

In the reference environment, proliferation rates decline with increasing distance from the tumor edge as a result of nutrient consumption of peripheral cells (Extended Data Fig. 5a). When peripheral cells die in response to therapy, resource availability in the tumor core improves, allowing previously quiescent cells to resume proliferation and contribute to the tumor regrowth. In addition, proliferating cells within the tumor bulk generate mechanical forces that push neighboring cells outward, leading to a cumulative collective displacement that increases towards the front (Extended Data Fig. 5b).

To augmented the training environment accordingly, we first introduce a distance-dependent growth rate for the resistant population, γ_*R*_(*d*), where *d* denotes the distance of a resistant clone to the tumor front (Fig. 5a). Second, we abstract the mechanically induced outward motion of resistant clones by introducing a distance-dependent velocity term, *v*(*d*), that governs the clone’s radial displacement toward the tumor edge. The resulting dynamics of the resistant population are described by the following system of equations:

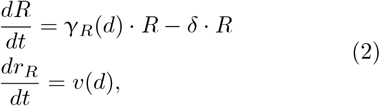

with *d* = *r*_*T*_ − *r*_*R*_, where *r*_*R*_ is radial position of the resistant clone and the tumor radius is approximated as 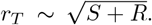 Both γ_*R*_(*d*) and *v*(*d*) are modeled as quadratically decaying functions of the form *f* (*d*) = *b*·(*a − d*)^2^, where parameters *a* and *b* are fitted to the position-dependent growth and velocity values measured in the reference environment (see Methods for details). Notice that the position-dependent growth rate in the augmented environment effectively replaces the role of the carrying capacity in the original base training environment.

We test the effects of these augmentations by evaluating the original agents — trained in TE_0_ — in the augmented environment and find that individual agent performance in the augmented environment correlates strongly with their performance in the reference environment (Extended Data Fig. 6a). Interestingly, both reference and augmented environments show a biphasic growth pattern that is in line with reports from previous *in vitro* studies (Fig. 5b)[26].

Finally, we train a new cohort of agents in the augmented training environment (Fig. 5c,d) (Extended Data Fig. 6b, Supplementary Movie 7). Cross-environment evaluation via GRAPE now show that these second-generation agents not only converge during training but now also exhibit superior and consistent performance in the reference environment (Fig. 5e).

Analyzing the strategies of these second-generation agent, we find that they consistently avoid treatment at low tumor burden, precluding the ratchet-like forward motion of resistant cells is prevented (Fig. 5c, d; Extended Data Fig. 7). Instead we observe a stable oscillation in clone distance (Fig. 5f, g). This success is further confirmed by the distribution of resistant cells remaining narrowly confined within the population bulk (Fig. 5h).

In addition to uncovering the importance of physical mechanisms for therapy dynamics, our findings point out a potential pitfall for the future design of adaptive therapy trials. An important insight from the adaptive therapy pilot study was the large variation of model parameters obtained from post hoc fitting [2, 3]. To more finely tailor therapy to individual patients, future trials might use an initial “probing cycle” for patient-specific model parameterization [46]. In such a scenario, patients would constitutively receive an initial round of fixed-threshold-therapy (e.g. AT_50_) to gauge the patient-specific tumor response for model parameterization. Tailored models can then be used to suggest optimal personalized scheduling, including RL-based approaches.

However, our findings suggest that such a probing cycle should be applied very carefully, as the associated decrease in tumor burden carries the risk of an unrecoverable competitive release of a pre-existing resistant clone. Fig. 5i shows a trajectory for an initial probing cycle with a 50% reduction in burden, followed by optimal RL-based therapy decisions (TTF = 26 ± 14 generations, *n* = 50). To mitigate the risk, data acquisition for parameterization could be split across multiple cycles of a higher-threshold-strategy (e.g. AT_75_) substantially lowering the probability of premature competitive release (Fig. 5j; TTF= 147 *±* 52 generations, *n* = 50).

## Discussion

In this work, we introduce Reinforcement Failing, a framework for the AI-guided, human-in-the-loop refinement of abstracted training environments for complex dynamics, such as tumor therapy. Using cross-environment group-relative agent performance evaluation, Reinforcement Failing helps to expose previously hidden emergent processes that impact therapeutic outcomes. This approach extends the application of RL beyond classic policy optimization toward artificial scientific discovery, offering a systematic way to identify and integrate emergent phenomena into interpretable training environments for enhanced transferability.

Applied to tumor therapy, the framework reveals that position-dependent growth and motion of resistant cells can drive a ratchet-like conveyance of resistant clones to the tumor front. As a result, even short treatment pulses can lead to irreversible loss of resistance control. Physics-informed environment augmentation enables agents to avoid these pitfalls and achieve superior therapeutic performance. Finally, our findings highlighting potential risks of ill-timed probing cycles for patient-specific model parameterization and outline mitigation strategies during translation.

Our findings add a new layer to recent advances in describing the population dynamics of densely packed cellular population, such as tissues and microbial biofilms, as emergent phenomena in actively proliferating granular matter[24, 25, 30, 47]. By tying these phenomena directly to adaptive tumor therapies via RL policy failures, we offer a structured way to incorporate these features into interpretable mathematical models of tumor therapy dynamics — a process that resonates with broader efforts in multi-fidelity and group reinforcement learning to overcome the limitations of “one-size-fits-all” training environments [35, 36, 48].

As with any modeling framework, Reinforcement Failing has inherent limitations that warrant careful consideration. First, detecting missing features depends on sufficient diversity among RL agents; without carefully designed reward functions and training initiation, the ensemble may converge to near-identical strategies, limiting our ability to glean insights on the wider behavior space structure. Second, although we employed a state-of-the-art physics-driven cancer simulator (PhysiCell), real tumors involve many additional factors such as vascular heterogeneity, immune infiltration, and metabolic feedback loops. Third, in a tumor context, the initial positioning and abundance of resistant cells in our reference environment, as well as the assumption of an isolated lesion, does not reflect the complexity encountered clinically [33]. The presented results should therefore be primarily interpreted as a conceptual demonstration of the Reinforcement Failing approach with a limited translational perspective. To overcome these limitations, future work will have to take these biological complexities into account while 11 pairing them with technical advances of RF, such as diversity-promoting mechanism during agent training.

One key avenue will be the integration of more physiologically detailed simulations that better incorporate features like angiogenesis or the mechanical tumor microenvironment. A particularly exciting avenue of research will be the integration of immunotherapy into the adaptive therapy framework[49].

Looking beyond oncology, employing Reinforcement Failing in other complex systems — such as ecological management[50], autonomous driving[51], or robotics[52] — could similarly reveal hidden emergent processes that hamper straightforward RL-based optimization.

On the artificial scientific discovery level, refining methods that intentionally promote policy diversity during training will help to ensure that the group-level performance analysis yields rich insights into underexplored parts of the decision space. On the long run, combining RL-guided mechanistic discovery with reasoning Large Language Models, akin to the recently introduced algorithmic discovery by AlphaEvolve[53], has the potential to extend exploration to realms not attainable by conventional research.

In summary, Reinforcement Failing reframes the perceived “failure” of RL policies to generalize as a potent discovery mechanism for uncovering cryptic emergent phenomena in high-dimensional, physics-driven environments. By systematically identifying and integrating missing processes, researchers can iteratively refine their mechanistic models and develop more robust, interpretable strategies for disease management. This work thus bridges the gap between realistic tumor behavior and simplified mathematical abstractions, laying a foundation for the RL-guided development of more nuanced therapy strategies. Ultimately, Reinforcement Failing offers a versatile tool for artificial scientific discovery in other domains as well, underscoring the value of embracing failure to achieve deeper insights.

## Methods

### Reference environment

The reference environment is modeled using an off-lattice agent-based cancer model (ABM) implemented in PhysiCell (v1.12.0) [38] with BioFVM (v1.1.6) [54], an open-source state-of-the-art cancer simulation platform [39–42]. The model captures cellular growth, physical interactions, and the diffusion of resources and therapeutics within a 2D domain of 2 × 2 mm, in which individual cells have a radius of 8.41 µm.

Cell proliferation follows a single-phase cycle regulated by oxygen availability, which diffuses from Dirichlet boundary nodes arranged in a circular pattern near the simulation domain’s perimeter. Cellular oxygen consumption establishes a radial gradient, restricting proliferation to an outer growth layer (Extended Data Fig. 5e-g), beyond which cells become quiescent. The parameters governing this oxygen-dependent behavior are informed by experimental data from Grimes et al. [55]. A full list of parameters is provided in the shared repository.

Cell mechanics and motion are governed by the inertialess equation of motion 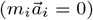, as described in Ghaffarizadeh *et al*. [38]. Adhesion and repulsion potentials are modeled using distant-dependent potential functions [54, 56]:

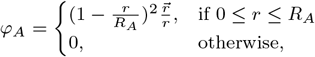

where φ_*A*_ denotes the adhesion potential, *r* is the distance between two cells, and 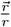 is the unit vector from one cell to another, and *R*_*A*_ is the maximum adhesive interaction distance.

Similarly, the repulsion potential is defined as:

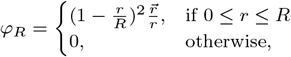

where φ_*R*_ represents the repulsion potential and *R* is the cell radius.

The initial tumor environment is generated by growing a tumor from a single cell over 15 generations (Supplementary Movie 1), resulting in a population of 3584 cells. We then locate a compact subpopulation of 5 drug-resistant cells approximately halfway between the tumor center and its surface.

Therapy is simulated by binary switching of Dirichlet boundary conditions, toggling between drug absence and full-dose exposure. Drug effects scales linearly with its concentration, largely increasing the apoptosis rate in proliferating cells. Cells growing slower than 10% of the maximum speed are not affected by the drug. Given the oxygen-dependent proliferation pattern, this results in preferential cell kill at the tumor periphery, where growth and drug concentrations are highest. In the absence of treatment, the baseline cellular turnover rate remains at 1%.

To apply RL policies we wrapped the aforementioned PhysiCell simulations in a custom Gymnasium environment [57]. Communication between the RL environment and the simulation was implemented using ZeroMQ [58], employing local inter-process communication (IPC) on Unix systems and Transmission Control Protocol (TCP) on Windows (see github repository for a full description).

### Base training environment

For the computationally efficient base training environment, we employ a competition model based on the system of two ordinary differential equations (1), previously well-studied in the context of adaptive therapy [20, 30, 43].

Populations of sensitive *S* and resistant *R* cells grow with rates γ_*S*_ and γ_*R*_. Their effective proliferation is regulated by the extend to which the total population *S* + *R* approaches the carrying capacity *K*. To capture the competitive disadvantage of resistant cells in the early phase of treatment, we introduce an asymmetric competition term *c*, which, if greater than 1, strengthens the suppressive effect of *S* on *R*. This adjustment is necessary to align the simplified model with the resistant suppression dynamics observed in the reference environment during reference AT_50_ therapy.

Both populations experience a constant death rate δ, independent of carrying capacity. Treatment is modeled as a fixed depletion *D* of sensitive cells, representing a constant number of cells reached and killed by the drug. This approximation reflects cell death dynamics in the reference environment when sensitive cells dominate and resistant fraction remains low. The fixed-cell-number killing term is motivated by resource-limited therapy: only cells within the well-perfused growth layer receive sufficient oxygen and drug exposure to be eliminated. In the tumor size regime relevant for this study, this approximation is justified by the approximately constant thickness of the growth layer (see Extended Data Fig. 5e-g). It should be noted, however, that neither this nor the more common proportional-kill formulation fully captures sensitive-cell death dynamics simultaneously across all scenarios, particularly under continuous or adaptive therapy (Fig. 1f,g).

Deviations between the non-spatial base and the spatial reference environments emerge in two key scenarios:

1. Small total population size, where a larger fraction of cells exhibit high growth rates, leading to higher-than-expected drug-induced death.
2. High resistant cell burden, which reduces the number of drug-exposed sensitive cells due to spatial constraints.

These discrepancies suggest potential refinements to the mechanistic mathematical model, incorporating spatial constraints and heterogeneous drug exposure.

### Calibration of the base training environment

We performed Bayesian inference to parametrize the base training environment using the PyMC package v5.9.2 [59]. Markov Chain Monte Carlo (MCMC) sampling was conducted with the Differential Evolution Metropolis (DEMetropolis) algorithm, running 8 parallel chains to ensure robust posterior estimation.

The carrying capacity *K* was fixed at 18,000 cells, corresponding to the estimated total cell count within the full simulation space. Parameter fitting followed an iterative process to ensure convergence and stability of parameter estimates:

1. The resistant cell growth rate γ_*R*_ was first fitted to a continuous therapy scenario, keeping all other parameters fixed.
2. The sensitive cell growth rate γ_*S*_, treatment strength *D*, and competition coefficient *c* were then fitted to the adaptive therapy (AT_50_) reference scenario, using the updated γ_*R*_.
3. This process was repeated iteratively until parameter changes between successive iterations were below 1%, ensuring model stability and accuracy in representing tumor dynamics across both reference therapy regimes.

For the final inference step, we used 2000 tuning steps followed by 1000 posterior draws. Priors were initialized as truncated Gaussian distributions, with mean values guided by previous analyses. Since priors evolved iteratively, we report the final prior means used in the last fitting iteration, which also represent the inferred parameter values used to train the RL agents: γ_*S*_ = 0.04, γ_*R*_ = 0.134, *D* = 330, *c* = 4.78. The inferred parameter values indicate a strong fitness advantage of resistant cells, while also highlighting a significant suppressive effect of sensitive cells on resistant cell growth.

For continuous (CT) and adaptive (AT_50_) therapies, treatment decisions were made every quarter generation to maintain cell counts near the treatment thresholds. For RL-based agents, this decision frequency was reduced to 1.5 simulated generations to increase clinical feasibility.

### Therapy performance metrics

As a robust metric for therapy performance, we measure the time to therapy failure (TTF), which we define as the time at which resistant cells reach the initial tumor size. TTF is analogous to the commonly used time to progression (TTP), defined via a fixed threshold on total population size relative to the baseline burden. TTF more robustly predicts the potential impact of continued treatment. For example, it precludes the previously reported “premature progression” scenario, in which a trajectory is falsely defined as progressed despite low resistance prevalence, due to a temporary fluctuation in tumor burden[46]. It should be noted, however, that the use of TTF is exclusive to *in silico* scenarios, where the abundance of resistant cells is readily available.

### Reinforcement learning

Treatment decision agents were implemented using Stable Baselines3 [60], a PyTorch-based library providing reliable RL algorithm implementations. We trained the agents using Proximal Policy Optimization (PPO) [61], with identical policy and value networks — both simple MLPs with two hidden layers of 128 units.

To account for history dependence and enable learning beyond simple threshold-based treatment rules, the observation space included stacked past 64 cell counts and treatment decisions, concatenated into a single observation. Hyperparameters (e.g., budget, entropy coefficient, clip range, batch size) remained consistent across agents and are available in the shared code repository.

Agents were trained for 5 × 10^6^ environmental steps and underwent ∼ 4, 900 policy update iterations, experiencing approximately 10^5^ full episodes throughout training. The reward function was designed to maximize time to progression while preventing excessive tumor burden. Specifically, agents received a reward of +1 per time step if the average tumor size remained below the initial level:

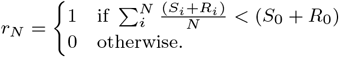

The base mathematical model and augmented mechanistic model were implemented in a gymnasium environment and solved using Euler’s method for training. To enhance agents’ policies robustness, Gaussian noise of mean 0 and *σ* = 0.01 · *N* was added to the calculated cell number *N* for both resistant and sensitive populations after solving the equations.

To assess the strength of the relationship between agent policy performance and the smallest tumor size at which each agent initiated treatment (Fig. 3d), we computed Pearson’s correlation coefficient (*r*) using the NumPy function corrcoef.

On 2 cores of Intel Xeon IceLake Platinum 8360Y processors, training times per agent were approximately 1 hr 58 min in TE_0_ and 2 hr 14 min in TE_+_ per agent.

### Group relative agent performance evaluation (GRAPE)

GRAPE is a performance evaluation framework based on relative cross-environment validation of agents. A cohort of equivalent agents (differing only in weight initialization) is first trained, and their performance is assessed in both the training and reference environments. This shifts the RL objective from identifying a single best-performing agent to exploring a broader decision space and enabling more reliable policy transfer to complex settings.

Cross-environment performance is quantified using a cross-environment performance difference (CEPD) map, which measures the standardized mean difference (Cohen’s *d*) between reference and training environments for each agent:

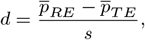

where 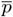 is the average performance of the agent and *s* is the pooled standard deviation.

In addition, GRAPE incorporates Kullback–Leibler (KL) divergence to compare policies directly. KL divergence quantifies the behavioral distance between agents, allowing us to determine whether policies achieving similar performance in TE rely on different decision strategies, and to visualize how performance differences between environments shift with changes in agent behavior.

Together, these measures reveal agents with strong or weak cross-environment generalization and help identify decision patterns contributing to poor transfer performance.

### Augmented training environment

To incorporate spatial dynamics into the mechanistic model, we modified the base ODE system (1). The sensitive cell population remained governed by its original equation (1a), while the resistant cell dynamics were updated with a position-dependent growth function (2).

Since oxygen availability defines cell growth rates in the reference environment, we approximated this dependence using distance from the tumor front — a reasonable proxy given that growth layer width remains relatively stable in large colonies (Extended Data Fig. 5e-g). We fitted this growth-distance relationship using free-growth simulations, excluding treatment cycles to avoid artifacts from loss of colony integrity. Similarly, we determined radially outward velocity as a function of cell position.

For position-dependent simulations, we estimated the total tumor radius (*r*_T_) based on total population size, assuming a densely packed tumor structure with cells of radius *r*_c_:

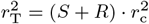

The resistant population was modeled using an effective radial coordinate *r*_*R*_, evolving over time as:

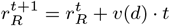

where velocity depends on distance to tumor front *v* = *f* (*d*).

Position-dependence of the growth rate *γ*_*R*_ and of the velocity *v* both have the form:

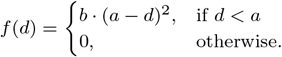

with *a* and *b* being fit parameters. Fits to the data from reference environment simulations are presented in Extended Data Fig. 5c,d.

## Data availability

Scripts and data used to produce the figures together with the trained networks are available at https://github.com/KayserLab/Aif_Reinforcement_Failing. Reference environment data, including cell positions, are available upon request due to their large size. Patient data from Ref. [3] used in Fig. 1 is available on GitHub (https://github.com/cunninghamjj/Evolution-based-mathematical-models-significantly-prolong-response-to-Abiraterone-in-mCRPC).

## Code availability

The code used for simulations, training, and analysis in this study is available at https://github.com/KayserLab/PhysiLearning. Scripts for plotting are available at https://github.com/KayserLab/Aif_Reinforcement_Failing.

## Acknowledgements

This work was supported by the Emmy Noether Programme of the German Research Foundation (project 455449456). ME acknowledges support from the German Academic Scholarship Foundation. The authors thank M. Dauber and L. Strampe for valuable discussions, I. Schneider for reviewing the manuscript, and J. Guck and his group for their invaluable support. We also acknowledge the open-source community, especially the contributors to Stable-Baselines3 and PhysiCell.

## Author contributions

S.A., and J.K. conceived and designed the study. S.A. performed the simulations and data analysis. S.A. and J.K. wrote the manuscript. S.A., M.E., N.A., E.F., T.C. and J.K. discussed and interpreted the results, prepared figures and contributed to the final revision of the manuscript.

---

**Extended Data Figure 1:**
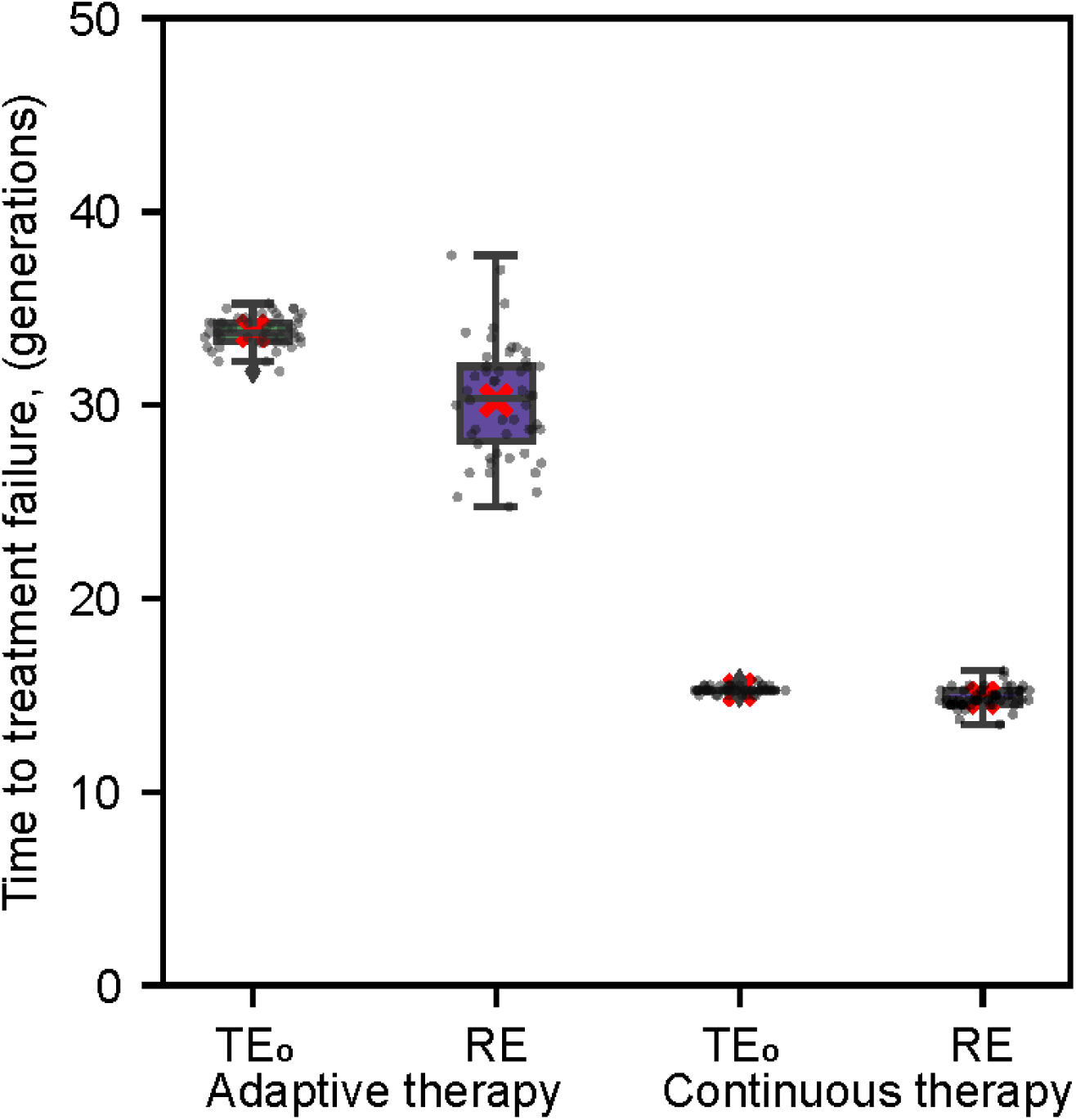
Reference therapies: Continuous Therapy (CT) and Adaptive Therapy (AT_50_ in base (TE_0_) and reference (RE) environments. Figure shows the time to treatment failure (TTF) distribution for the CT and AT_50_ therapy protocols applied to both the base and reference environments. Individual dots represent individual simulation runs, while the boxplot displays the median and quartiles. The whiskers extend to the rest of the distribution, and the red cross indicates the mean.

**Extended Data Figure 2:**
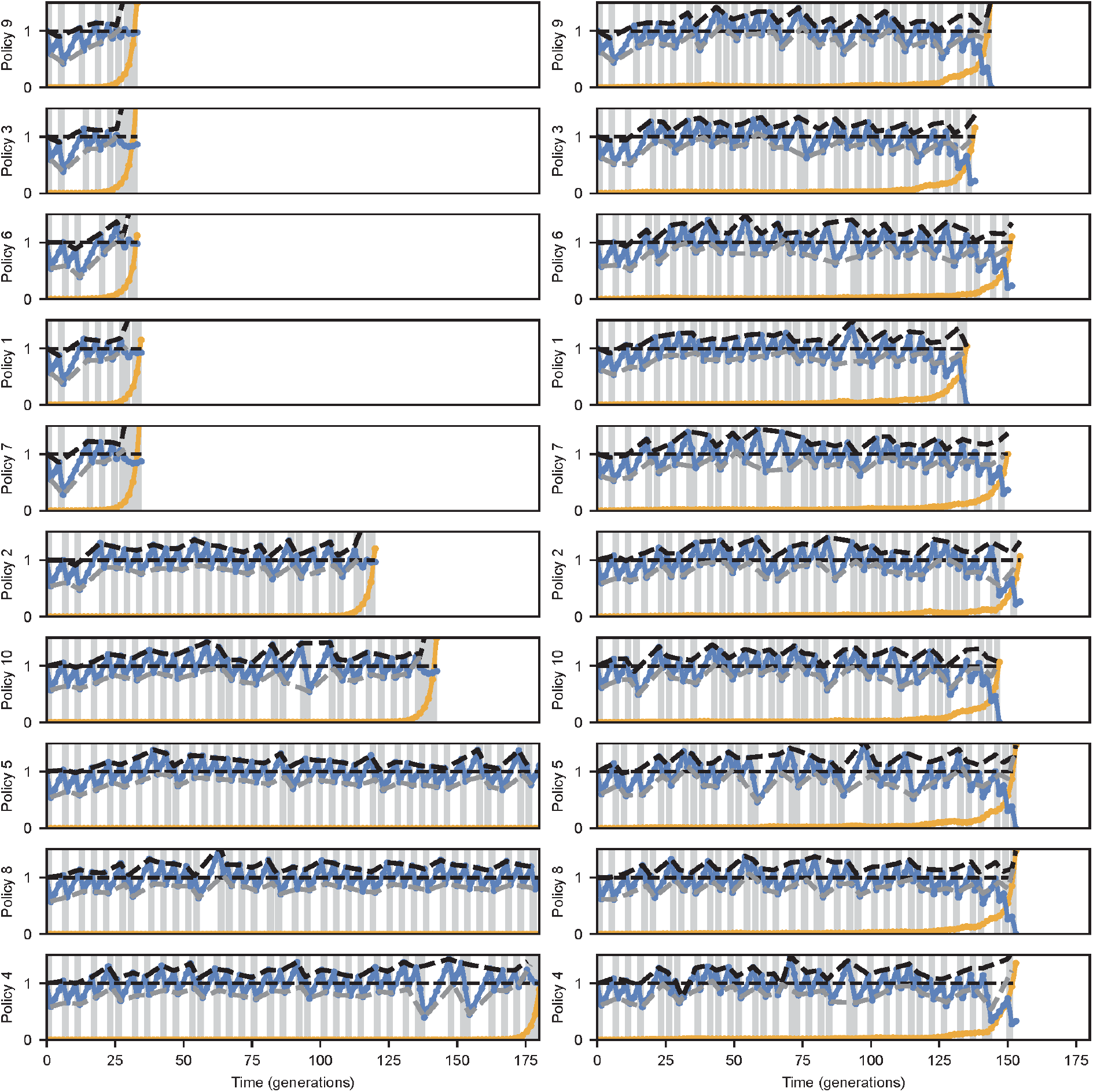
Sensitive and resistant population dynamics under agents policies trained in the base environment. Dynamics of the sensitive (blue) and resistant (orange) population sizes - relative to the total initial population size - under all n=10 policies, which were trained in the base environment (TE_0_). On the left - evaluation runs in the reference environment (RE), on the right - evaluation runs in the base training environment (TE_0_). Grey regions indicate treatment application. Dashed lines show the tumor burdens at treatment initiation (black) and treatment pause (grey). Each run corresponds to a median time to failure of the evaluated policy.

**Extended Data Figure 3:**
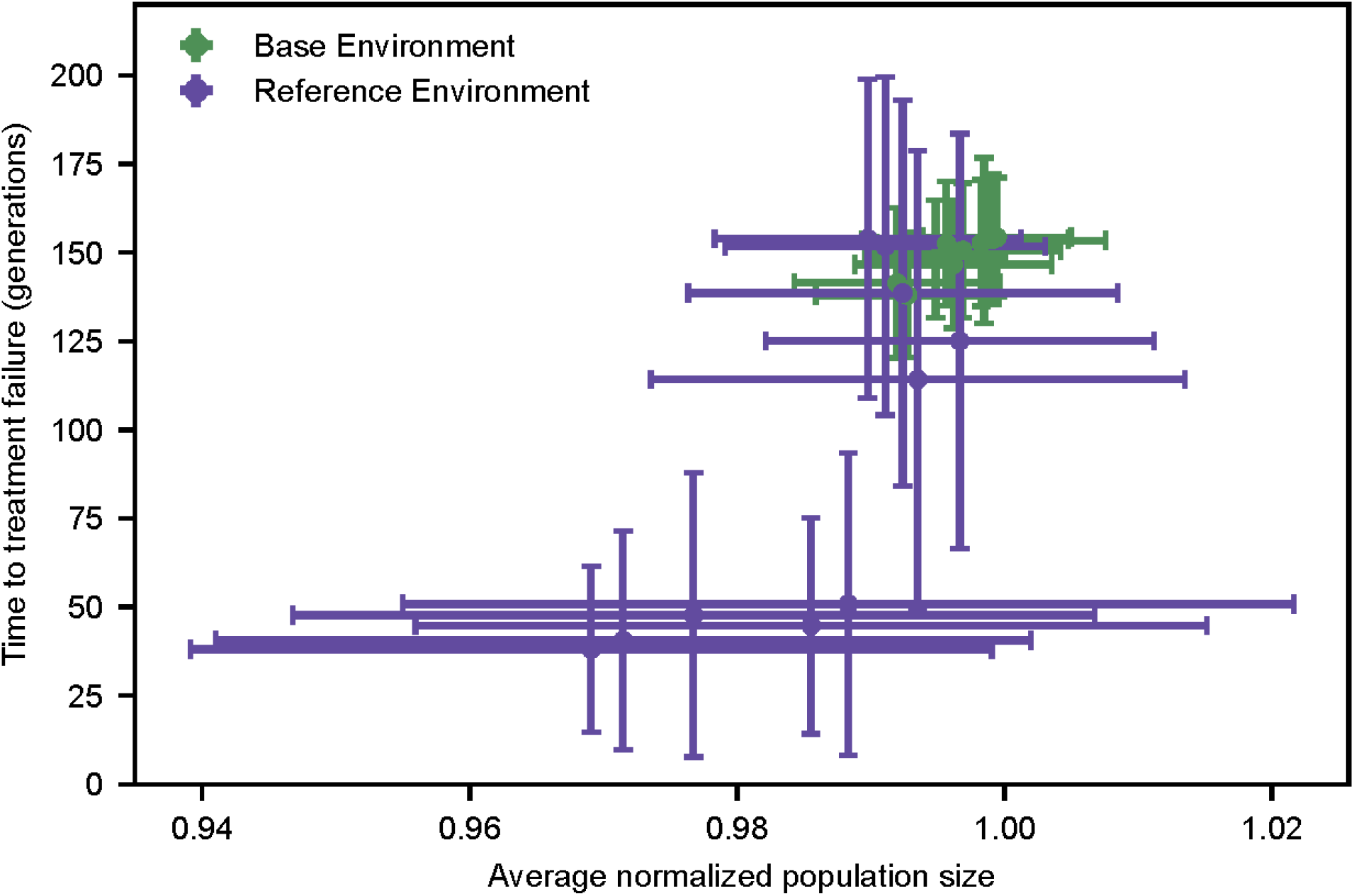
Time to treatment failure (TTF) in TE_0_ (green) and RE (violet) for n=10 policies trained in TE_0_ as the function of the average total population size. Dots represent averages over n=50 runs, and error bars indicate the standard deviation.

**Extended Data Figure 4:**
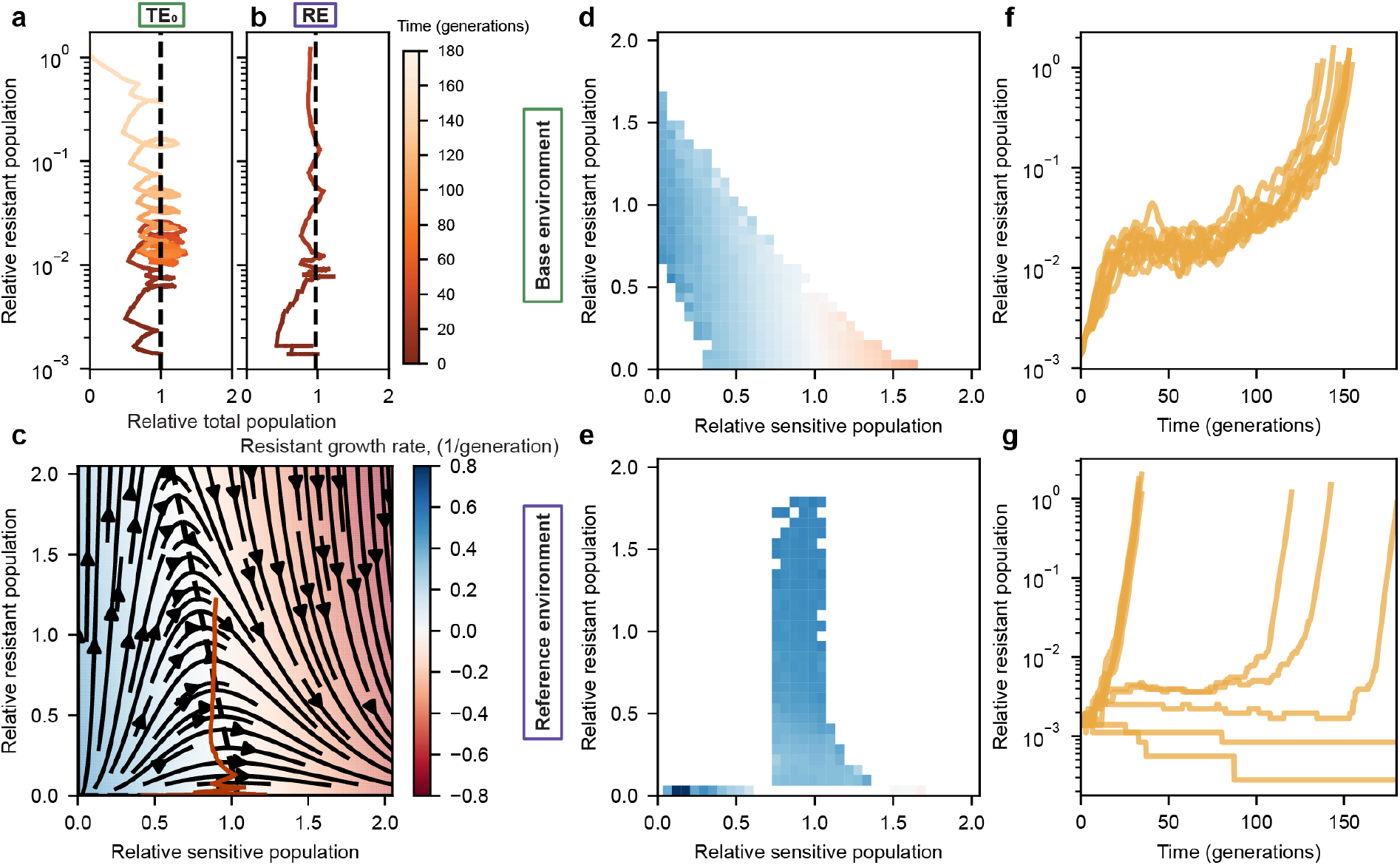
Control of resistant population in the reference environment is not governed by sensitive cell numbers. **a, b** Relationship between resistant and total population sizes over time (color) for the base (TE_0_) and reference (RE) environments during therapy under the worst policy. Dashed lines indicate the average cell number kept by the policy. **c** Phase space of resistant vs. sensitive populations in TE_0_. Streamlines represent trajectories in time under free growth, orange line shows the trajectory of the therapy run in RE, presented in **b**. Colors indicate the effective growth rate of the resistant population per generation. **d, f** Regions of the phase space explored by the n=10 agents over n=50 evaluation runs on TE_0_ and RE. The color scale from **c** is shared by **d** and **f. e, g** Resistant population dynamics over time for the evaluation run corresponding to the median TTF across all n=10 agents for TE_0_ and RE. All population sizes are shown relative to the initial tumor burden.

**Extended Data Figure 5:**
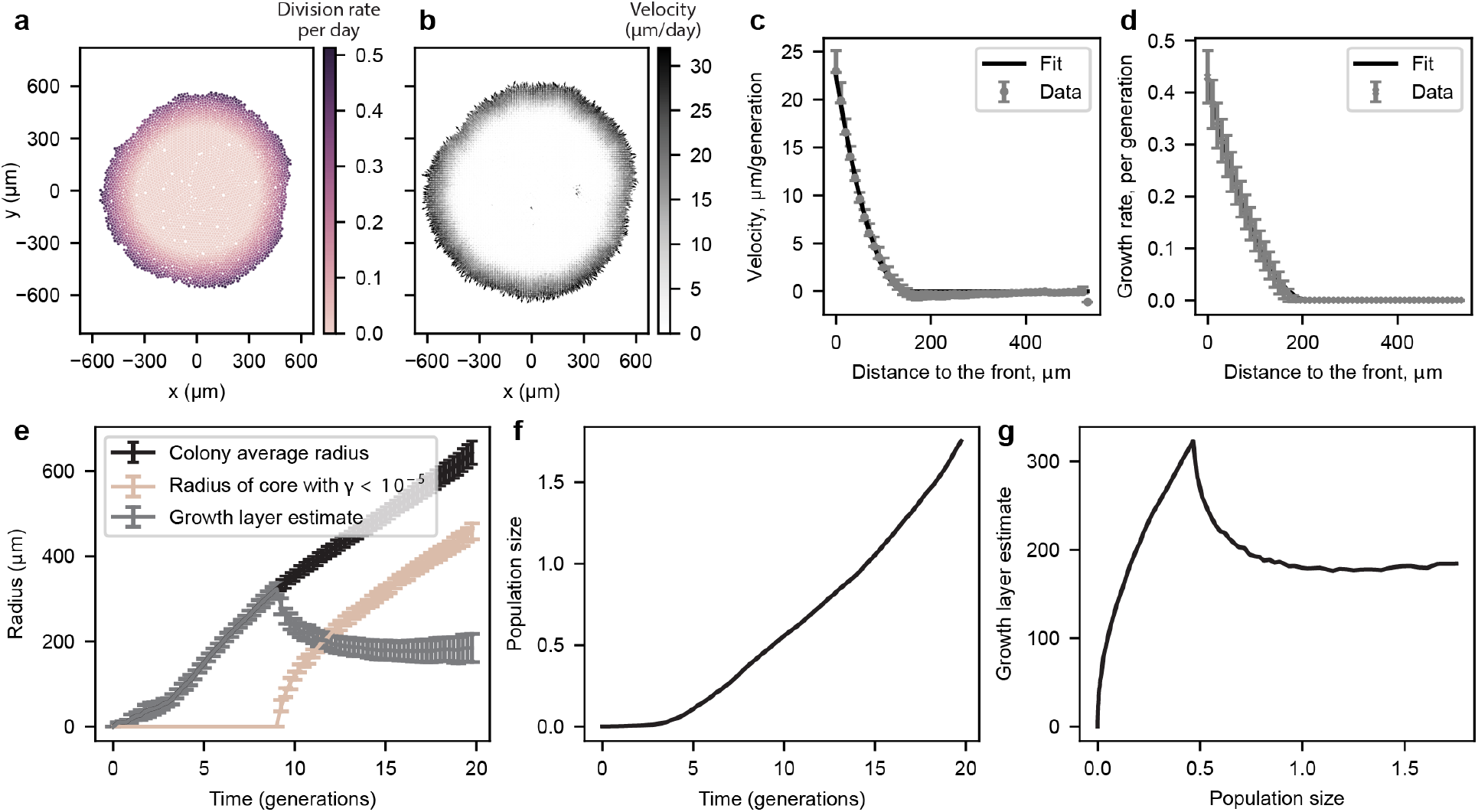
Position-dependent growth and motion of the cells. **a** Single-time snapshot of the 2-D growth rate distribution, where brightness decreases with growth rate. **b** Average velocity of cells at certain positions over 3 generations under free growth (n=50 runs). Arrow magnitude is proportional to velocity. **c, d** Fit of position dependent velocity and growth rate for the augmented environment. Data from the reference environment averaged over n=50 simulation runs over 3 generations of free growth is shown in grey. Solid black line represents the fit of a truncated quadratic function *f* (*d*) = *b*(*a − d*)^2^. **c** Error bars represent 25th and 75th quantiles. **d** Error bars show standard deviations. **e** Dynamics of the estimated tumor radius (black), the radius of the non-growing (*γ <* 10^−5^) core inside the tumor (beige), and the growth layer (grey), defined as the difference between the two, over time. The simulation is performed from a single cell seeded in the center of the simulation space over 20 generations. **f** Dynamics of the total population size, normalized to the initial population size used in training. **g** Relationship between the growth layer width and the relative population size.

**Extended Data Figure 6:**
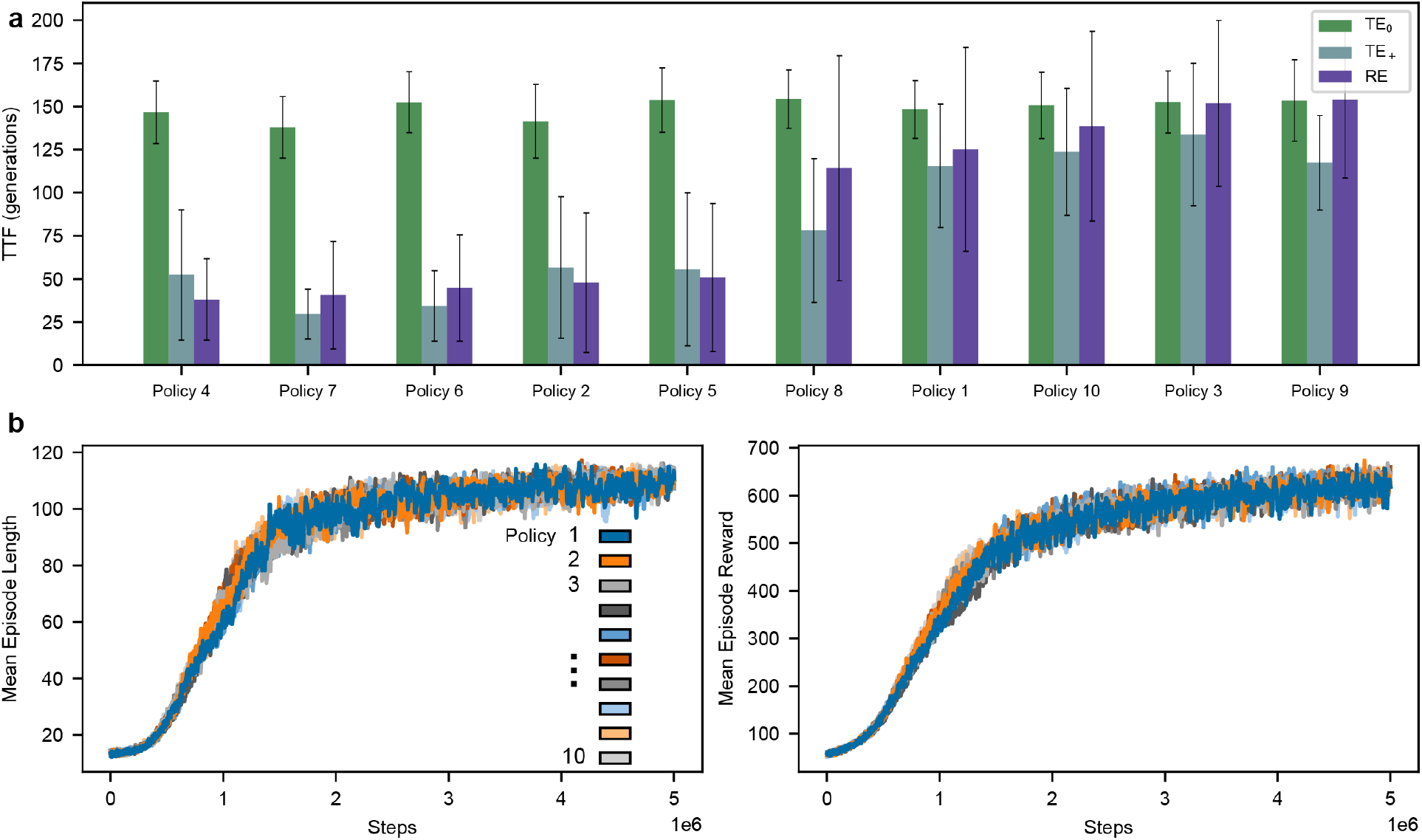
**a** Average time to treatment failure (TTF) achieved by the RL agent with policies trained in TE_0_, evaluated in TE_0_ (green), TE_+_ (blue-green) and RE (violet). n=50 runs, error bars indicate the standard deviation. **b** Mean episode length and mean episode reward for each individual training run in the augmented environment as a function of simulated steps. One steps represents one treatment decision point and corresponds to 1.5 simulated generations. Policy updates are performed every 1024 steps.

**Extended Data Figure 7:**
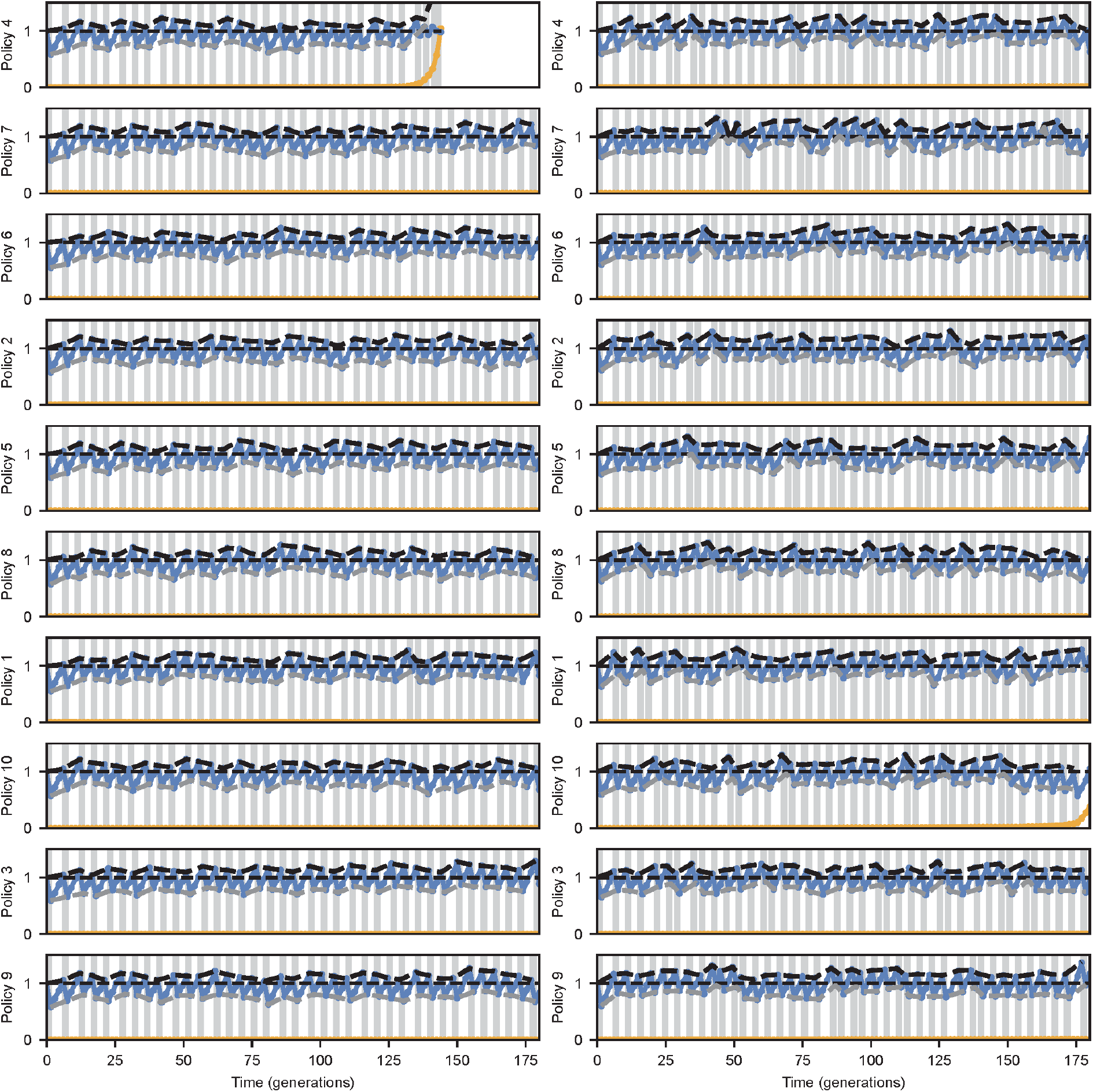
Sensitive and resistant population dynamics under agents policies trained in the augmented environment. Dynamics of the sensitive (blue) and resistant (orange) population sizes - relative to the total initial population size - under all n=10 policies, which were trained in the augmented training environment (TE_+_). On the left - evaluation runs in the reference environment (RE), on the right - evaluation runs in the augmented training environment (TE_+_). Grey regions indicate treatment application. Dashed lines show the tumor burdens at treatment initiation (black) and treatment pause (grey). Each run corresponds to a median time to failure of the evaluated policy.

