## Supplementary Text for "Reinforcement Failing guides the discovery of emergent physical dynamics in adaptive tumor therapy"

### 1 Supplementary Text: Control of resistant population is not governed by total number of sensitive cells in the reference environment

To investigate why early treatment at low tumor burden irreversibly leads to therapy failure in the reference environment, we analyze the relationship between resistant and total population sizes over time (Extended Data Fig. 4a-b). In the base training environment (Extended Data Fig. 4a), resistant cells initially expand rapidly during the early treatment phase when tumor burden is low. However, as the total population size increases later in the therapy cycle, resistant cell growth slows and can even reverse, leading to partial suppression. Between generations 30 and 80, therapy cycling maintains the resistant subpopulation in a controlled oscillatory state around 2%. While initiating treatment at lower tumor burdens in later stages weakens this control, resistance remains suppressed whenever the overall tumor burden is sufficiently high.

In contrast, the reference environment exhibits fundamentally different behavior (Extended Data Fig. 4b). Following aggressive early treatment, resistant cells initially expand at a high rate but are temporarily held in check. However, despite moderate tumor burdens in later treatment cycles, resistant cells eventually resume rapid growth with minimal suppression, even when total tumor burden remains high. The divergence between the two environments becomes particularly apparent when mapping population trajectories into a population phase space [1], using the relative sizes of the susceptible and resistant populations as dimensions (Extended Data Fig. 4c). In the training environment, trajectories follow the expected flow field from the underlying equations, where resistant cells can be re-suppressed after expansion. In contrast, the reference environment deviates sharply, suggesting that competitive release becomes effectively irreversible. These findings challenge a central assumption of the base environment — that resistance suppression is primarily dictated by high total cell numbers [2].

To further explore this discrepancy, we examine resistant cell growth under agent-driven therapies in the same phase space. In the training environment, observations align with model expectations, where resistance growth remains low or even negative when the susceptible population is large (Extended Data Fig. 4d). However, in the reference environment, the same phase-space region consistently exhibits strong resistant growth (Extended Data Fig. 4e), indicating that the expected suppression is absent. Temporal analysis of multiple resistance growth trajectories further highlights the contrast between environments: in the training model, growth follows predictable trajectories, while in the reference environment, resistance growth surges at widely varying time points (Extended Data Fig. 4f, g).

Together, these findings challenge the assumption that resistance suppression is primarily governed by the average abundance of susceptible cells. Instead, the loss of resistance confinement appears to be triggered by a transient reduction in tumor burden rather than absolute population numbers. In the next section, we will explore the underlying mechanisms driving these dynamics.
